# Convergent Roles of Growth Differentiation Factor-15 (GDF-15) in Mechanotransduction, Vascular Disorganization, and Immune Suppression in Melanoma

**DOI:** 10.64898/2026.01.14.699463

**Authors:** Yu-Chi Chen, Vishnu Sravan Bollu, Sina Kheirabadi, Kyle LaPenna, Arthur Berg, Ping He, Todd D. Schell, Amir Sheikhi, Erdem D. Tabdanov, Gavin P. Robertson

## Abstract

Melanoma, particularly in its advanced forms, remains one of the most lethal skin cancers, with limited effective treatments for both common cutaneous subtypes and rarer variants such as acral melanoma. The effects of the extracellular matrix (ECM) and other cancer-cell extrinsic processes on melanoma development remain underexplored. This study identifies Growth Differentiation Factor-15 (GDF-15) as a novel mechanosensing-regulated driver of melanoma pathogenesis across melanoma types. GDF-15 is shown here to be mechanically induced by ECM rigidity and compressive forces occurring during metastatic progression, leading to significantly elevated levels in both cutaneous and acral melanoma cells. In this study, ECM rigidity was recapitulated using cell-adhesive gelatin methacryloyl (GelMA) hydrogel microparticles (microgels) with tunable stiffness, providing a biomimetic platform to investigate how mechanical cues in the tumor microenvironment regulate GDF-15 expression in melanoma. A previously unrecognized synergy between GDF-15 and inflammatory factors commonly present in tumors was identified and found to promote a disorganized, hyperpermeable vasculature, thereby facilitating tumor nutrient access and impeding effective immune cell infiltration. GDF-15 knockdown reduced the intratumoral hemorrhage phenotype, indicating a causal role. GDF-15 also functions by directly suppressing natural killer (NK) cell-mediated cytotoxicity, revealing a second cooperating mechanism of immune evasion. These effects position GDF-15 as a key node linking mechanical stress, co-operation with inflammatory factors, abnormal vascular development, and immune dysfunction, thereby converging on a pathways and processes not previously described in melanomas.

**TEASER:** Uncovering GDF-15 as a central mediator at the intersection of mechanical rigidity and external force, disorganized vascular hemorrhagic remodeling, co-operation with inflammatory factors, and immune evasion, revealing a previously unrecognized therapeutic vulnerability across both common and rare melanoma subtypes

## INTRODUCTION

Melanoma is a highly aggressive skin cancer with a propensity for rapid progression, metastasis, and resistance to conventional therapies (1). Despite significant advances in early detection and treatment strategies, metastatic melanoma remains a major clinical challenge, with poor prognosis and limited therapeutic options (2). While immune checkpoint inhibitors and targeted therapies have revolutionized melanoma treatment, many patients experience resistance or relapse (3–5), highlighting the need to understand better the mechanisms driving tumor progression and immune evasion.

Tumor progression is influenced not only by genetic and epigenetic alterations but also by the interplay between cancer cells, the TME, and external factors (6,7). The TME, which includes blood vessels, fibroblasts, immune cells, stromal components, extracellular matrix (ECM), and signaling molecules, is pivotal in promoting tumor growth, angiogenesis, and immune suppression (6). Growth Differentiation Factor-15 (GDF-15), a member of the transforming growth factor-beta (TGF-β) superfamily (8), has emerged as a potentially critical mediator within the TME (9). Specifically, GDF-15 is a mechano-responsive cytokine that is upregulated in various cancers and contributes to tumorigenesis, metastasis, and immune modulation (10). Elevated serum GDF-15 levels have been associated with advanced disease stages and poorer clinical outcomes (11). GDF-15 is known to exert pleiotropic effects that enhance tumor cell survival and metastasis (8,9). Mechanistically, it has been implicated in promoting epithelial-to-mesenchymal transition (EMT) (12,13), modulating angiogenesis (14), and altering extracellular matrix dynamics (15). Additionally, GDF-15 may suppress immune function, thereby facilitating immune evasion (8).

Despite these observations, several critical gaps remain in our understanding of how GDF-15 contributes to melanoma progression. Most notably, the upstream biomechanical processes in tumors that regulate GDF-15 expression remain undefined. While GDF-15 is known to be induced by cellular stress, its regulation by structural ECM rigidity or mechanical cues within the TME remains unclear. Additionally, GDF-15’s interactions with inflammatory mediators and its influence on tumor architecture, including the development of disorganized vasculature and ECM, have not been reported. How these features collectively contribute to immune suppression and melanoma progression remains an open and essential question in the field.

Further complicating GDF-15’s role is the lack of a known receptor outside the central nervous system (16), and its association with immune suppression has primarily been attributed to limited T-cell infiltration (17,18). Its potential role in modulating innate immune responses, including the impairment of natural killer (NK) cell cytotoxicity, remains uncertain. Thus, a detailed understanding of how GDF-15 interfaces with the physical and immunologic components of the tumor microenvironment is lacking.

Given GDF-15’s multifaceted roles and clinical associations with poor outcomes (11), it has garnered interest as both a potential therapeutic target and a biomarker in cancer (8,19). However, its value in melanoma remains largely unexplored. Understanding the conditions that drive GDF-15 expression and the functional consequences for tumor architecture, angiogenesis, and immune surveillance could open new avenues for therapeutic intervention, particularly in immune-resistant or immune-excluded tumors.

This study addresses these key knowledge gaps by investigating the mechanistic underpinnings of GDF-15 regulation and function in melanoma. Specifically, we demonstrate that differential ECM rigidity, introduced via gelatin methacryloyl (GelMA) hydrogel microparticles (microgels) that partially mimic native tumor ECM, and applied physical pressure, mimicking the biomechanical stress observed outside of the cell during cancer progression, induces GDF-15 expression across melanoma types. Most significantly, we show that GDF-15 promotes tumor progression by synergizing with inflammatory factors in the TME, leading to vascular disorganization and immune suppression, particularly by impairing NK cell-mediated cytotoxicity. Finally, we confirm serum GDF-15 as a potential biomarker, based on tumor burden, and correlate its levels with the ECM rigidity and mechanical pressure experienced by tumor cells during cancer progression. Together, these findings establish GDF-15 as a central mediator at the intersection of mechanical rigidity and external force, disorganized vascular hemorrhagic remodeling, and immune evasion, and further support its status as a key target in melanoma.

## RESULTS

### Extracellular Matrix Stiffness and Mechanical Force Induce GDF-15 Expression

The TME in melanoma is increasingly recognized as a dynamic and mechanical ecosystem that profoundly influences disease progression (20), therapeutic resistance (21), and immune evasion (22). While much attention has been given to the deregulated biochemical signaling networks within the TME (23), the role of biophysical stress, particularly ECM rigidity and mechanical force, remains underexplored, especially in the context of melanoma subtypes such as acral melanoma, which arise in mechanically stressed anatomical sites on the palms of the hand and soles of the feet (24). Investigating whether GDF-15 is induced in melanoma cells in response to matrix rigidity and mechanical compression is critical to understanding how biomechanical cues shape tumor behavior, progression and immune interactions. If GDF-15 is a central mechanosensing-modulated effector in melanoma, it would represent a paradigm shift in how we conceptualize stromal-tumor communication, offering a novel mechanistic link between tissue biomechanics and melanoma aggressiveness.

In this context, we examined the impact of ECM rigidity and applied mechanical pressure on cell-mediated GDF-15 expression. **Figure 1A** provides an overview of the experimental model we have developed to demonstrate that GDF-15 can be induced in melanoma cells grown in culture and in animals *via* their mechanosensory function on the ECM, which varies in rigidity and mechanical force. We cultured melanoma cells with GelMA microgels with differing rigidity to form cell-microgel aggregates and then measured the levels of GDF-15 expression. Specifically, bulk hydrogel scaffolds were fabricated with varying GelMA concentrations (4 and 15% w/v), and their dynamic moduli, estimating the local rheological properties of corresponding microgels (25), were evaluated using rheological characterization, with each concentration corresponding to a different stiffness (**Figures 1B-D, Supplementary Figure S1**). The storage modulus for the soft and stiff scaffolds is 10.3 ± 1.5 kPa and 103.2 ± 10.6 kPa, and the corresponding loss modulus is 343 ± 59 Pa and 2850 ± 780 Pa, respectively (**Figures 1C-D**). Subsequently, GelMA droplets were fabricated using a high-throughput microfluidic device and characterized. The average sizes of the soft and stiff droplets were 31.8 ± 1.5 µm and 31.3 ± 1.7 µm, respectively (**Supplementary Figures S2A-C**). The droplets were initially stored at 4°C to facilitate physical crosslinking and were subsequently exposed to 395 nm light (2 W/cm²) for photo-crosslinking (**Figure 1E**). After oil and surfactant removal, the stability of microgels was assessed following a 60-min incubation at 37°C (**Figure 1F**). Their size distribution showed that the average diameters of soft and stiff microgels were 22 ± 1 and 26 ± 2 μm, respectively (**Figure 1G)** (25).

**Figure 1.**
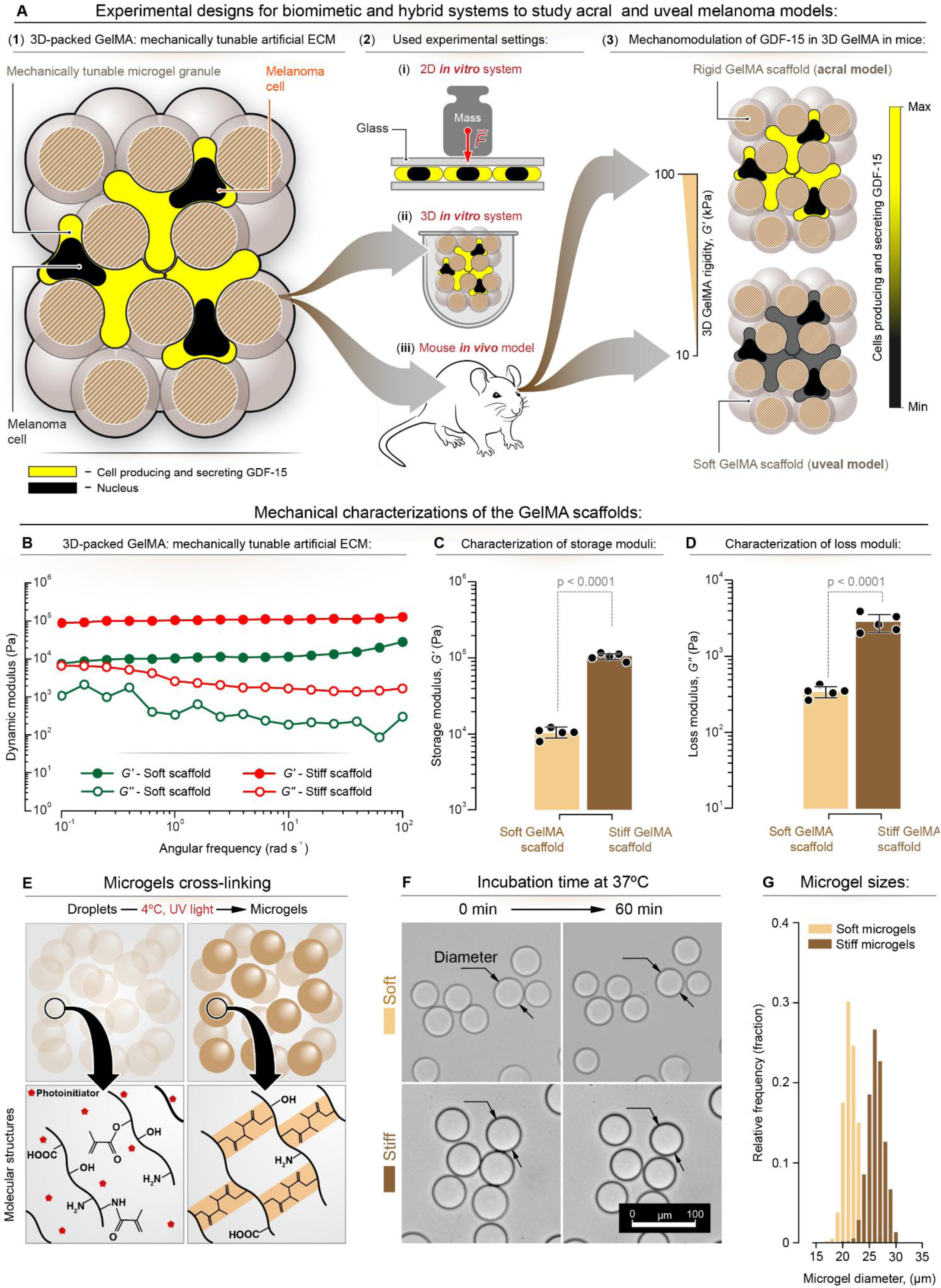
Schematic representation of the modelling used to show the effects of mechanical force and ECM rigidity on GDF-15 expression in melanoma cells. **(A)** Mechanical force exerted by applied weight on cells was used to examine pressure effects during metastatic processes. ECM stiffness was studied by comparing a rigid (*G*’ ≈ 100 kPa) environment resembling the acral melanoma model with a softer environment (G’ ≈ 10 kPa) in a GelMA hydrogel mimicking cutaneous melanoma. The color gradient (yellow to black) represents GDF-15 expression levels, with higher expression in the acral melanoma model. **(B)** Dynamic moduli (storage modulus, *G*′, and loss modulus, *G*″) of soft and stiff bulk hydrogel scaffolds, made with 4 and 15% w/v GelMA concentrations, respectively, versus the angular frequency at a constant oscillatory strain (∼ 0.1%). **(C)** *G*′ and **(D)** *G*″ of soft and stiff scaffolds at a constant angular frequency (1 rad s^-1^) and constant oscillation strain (0.1%) (n = 5). **(E)** GelMA droplets (4 or 15% w/v) in oil and surfactant are stored at 4 °C to create physically crosslinked microgels, followed by photo-crosslinking to fabricate thermally stable GelMA microgels. **(F)** Brightfield microscopy images of soft and stiff microgels incubated in 37°C DPBS for 60 min, imaged at 15-min intervals. Scale bar is 100 µm. **(G)** Size distribution of soft and stiff microgels after incubation at 37 °C for 60 min (*n* > 800). A one-way analysis of variance (ANOVA), followed by Tukey’s post hoc multiple comparisons test, was performed to compare the study groups.

To assess whether ECM rigidity-mediated induction of GDF-15 is directly proportional to ECM rigidity, melanoma cells were cultured on GelMA microgels of varying stiffness. Specifically, cutaneous UACC 903 and A375, and acral WM4235 melanoma cell lines were seeded with GelMA microgels with two distinct stiffness levels: soft (*G’ =* storage modulus ≈ 10 kPa) and stiff (*G*’ ≈ 100 kPa). Cells and GelMA microgels were co-seeded at a 1:3 ratio (cell: microgel) in low-adhesion round-bottom microwell plates and maintained for 24 h before staining for GDF-15, F-actin, and the Hoechst dye nuclear markers. Quantitative fluorescence imaging was used to assess a stiffness-dependent level of GDF-15 expression, showing that all melanoma cells exhibited an induction of GDF expression (**Figures 2A, B, C**) with those on stiff microgels displaying two to threefold higher expression (**Figure 2D**). For added experimental rigor, Western blot analysis of the cells showed increased GDF-15 expression relative to negative controls and comparable to that observed for the positive controls (**Figure 2E**). These findings underscore the role of ECM rigidity in regulating GDF-15 expression in melanoma cells.

**Figure 2.**
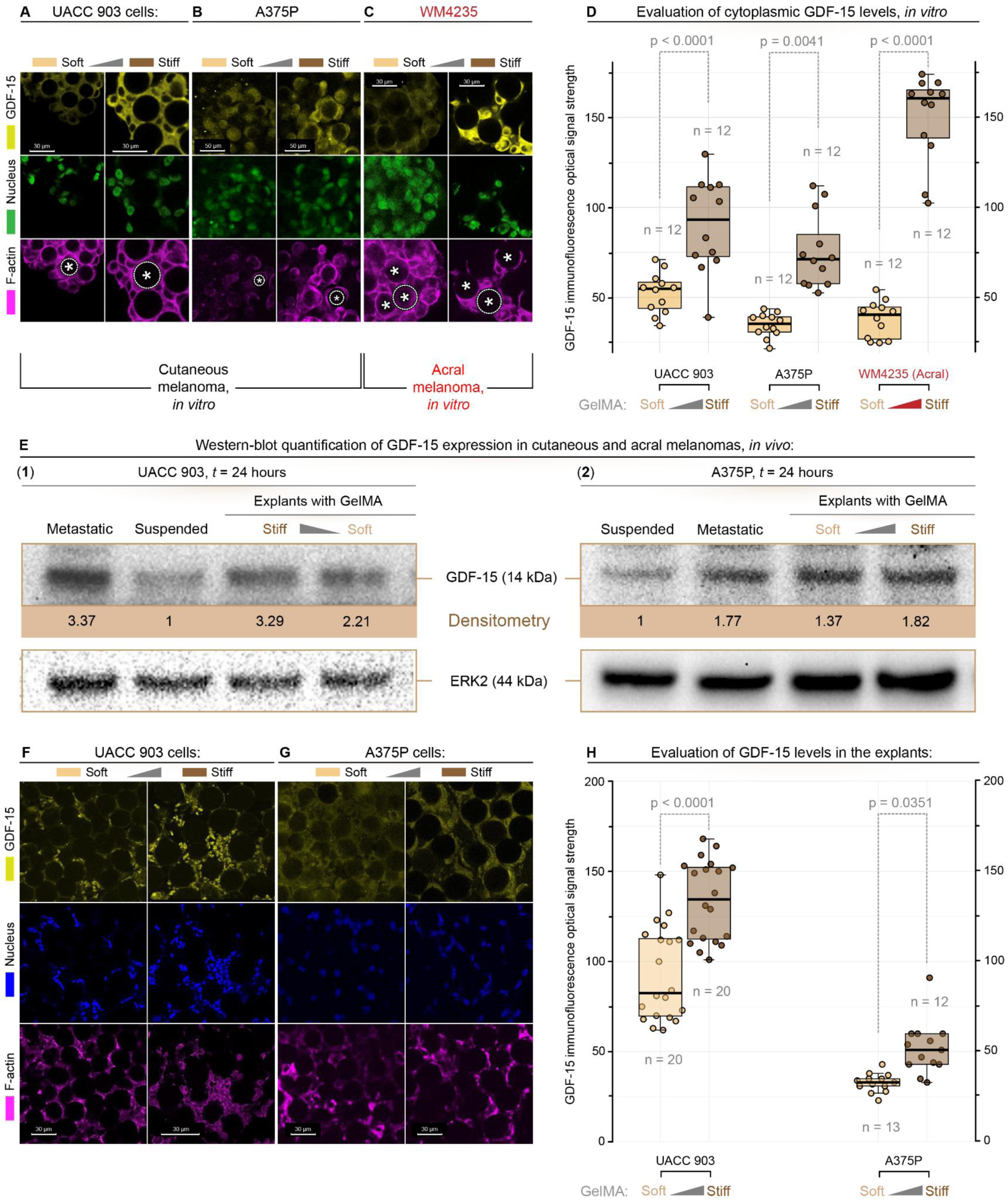
ECM rigidity modulates GDF-15 expression in melanoma cells cultured on GelMA_microgel. Representative immunofluorescence images showing GDF-15 expression (yellow), nuclei (green), and F-actin (magenta) in melanoma cells (**A**: UACC 903; **B**: A375P; **C**: WM4235) cultured with soft (*G*’ ≈ 10 kPa) and stiff (*G*’ ≈ 100 kPa) microgel matrices. Increased GDF-15 expression is observed in the stiff microenvironment, resembling conditions found in acral melanoma. **(D)** Quantification of GDF-15 fluorescence intensity in melanoma cells (Cutaneous: UACC 903 & A375P; Acral: WM4235) cultured in soft and stiff matrices. Box graphs represent median ± SEM. **(E)** Western blot analysis of GDF-15 expression in melanoma cells (UACC 903 & A375P) under soft or stiff GelMA ECM matrix conditions. ERK2 serves as a loading control, and GDF-15 protein levels are quantified and normalized to ERK2. Cells in the stiff matrix show increased GDF-15 expression compared to the soft matrix. (F, G, H) Modeling GelMA microgel stiffness in melanoma tumors. **(F, G)** Representative immunofluorescence images showing GDF-15 (yellow), nuclei (blue), and F-actin (magenta) in tumor sections (F: UACC 903; G: A375P) after 7 days of growth in mice. (H) Quantification of GDF-15 fluorescence intensity, equivalent to expression in tumor cells after growing on stiff or soft microgels.

To investigate GDF-15 induction in a similar animal model, a subcutaneous melanoma model was established using GelMA microgels. Melanoma cells were mixed with microgels at a 1:1 ratio (cell: microgel), allowed to attach and interact for 24 h, and subsequently injected subcutaneously into mice. Tumors were harvested seven days post-injection, sectioned, and immunostained for GDF-15 (**Figure 2F).** Representative immunofluorescence images showing GDF-15 (yellow), nuclei (blue), and F-actin (magenta) in tumor sections (**F**: UACC 903; **G**: A375P) after 7 days of growth in mice. Quantitative microscopy revealed a stiffness-dependent increase in GDF-15 expression for UACC 903 (**Figure 2F**) and A375P (**Figure 2G**) melanoma cells associated with rigid GelMA microgels exhibiting higher GDF-15 levels compared to those attached to soft microgels (**Figure 2H**). This observation reinforced the role of matrix stiffness in regulating GDF-15 expression in melanoma cells within the in vivo tumor microenvironment.

The effect of substrate type on GDF-15 expression was examined by culturing human UACC 903, and mouse B16-F10, and M4 (26) melanoma cells on plastic, collagen-coated surfaces, or in suspension (**Figure 3A**). Cells grown on collagen exhibited the highest GDF-15 induction, followed by those in suspension. UACC 903M cells constitutively overexpressing GDF-15 were used as a positive control. Mouse melanoma cells were included for rigor and to demonstrate similar effects across species. These results underscore the influence of both ECM rigidity and force on GDF-15 expression in melanoma cells. Additionally, the role of external mechanical force in GDF-15 induction was next evaluated using a weight-based compression assay model diagrammatically illustrated in **Figure 3B**. Human UACC 903 or mouse B16-F10 melanoma cells were grown on coverslips, inverted, and subjected to weights of 1 g, 2 g, or 5 g for 24 h. Western blot analysis revealed a dose-dependent increase in GDF-15 expression with increasing weight for human UACC 903 (**Figures 3C, D**) and mouse B16F10 (**Figures 3E, F**) melanoma cells.

**Figure 3.**
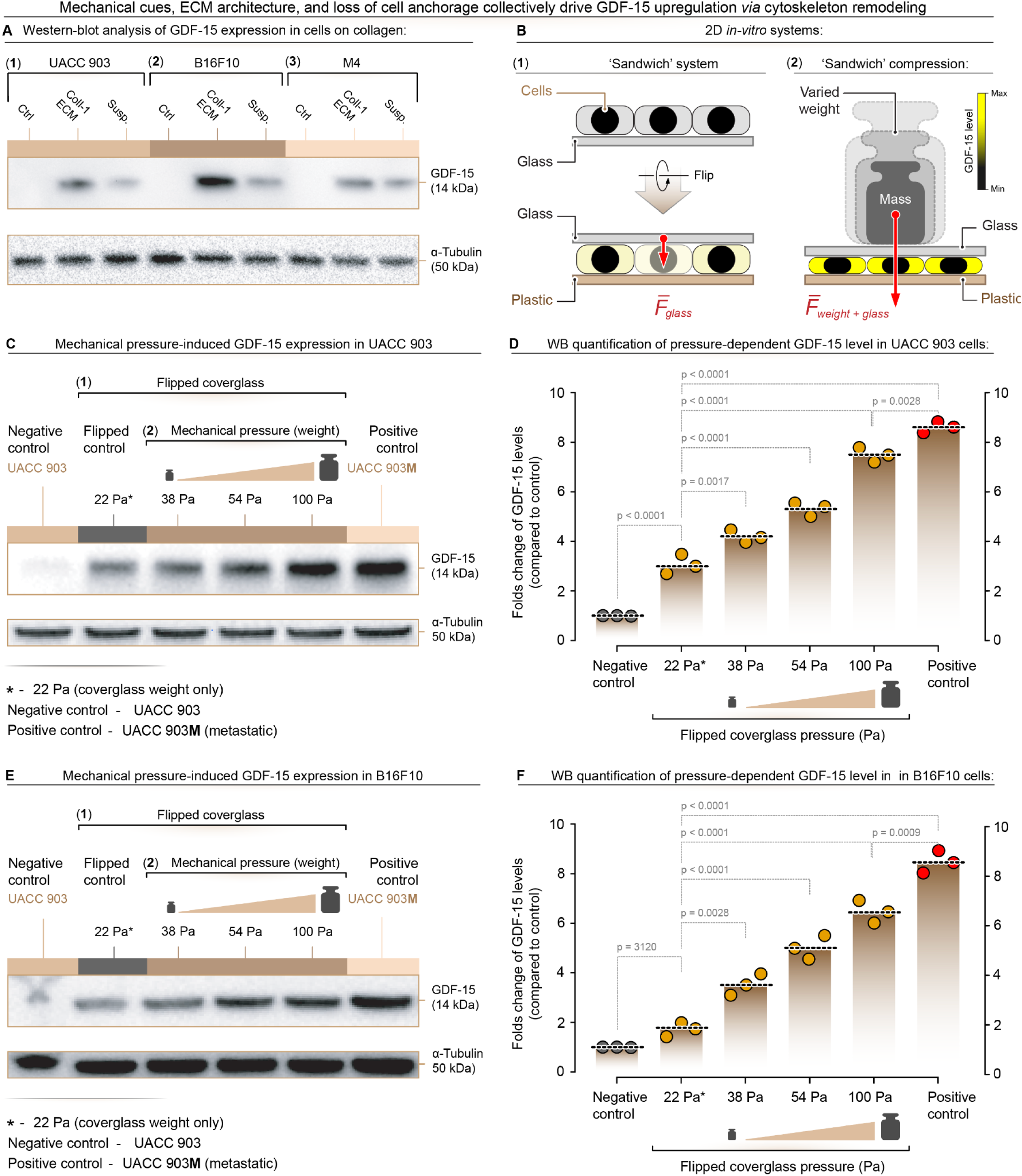
Mechanical force, ECM complexity, and anchorage independence regulate GDF-15 expression through cytoskeletal remodeling. **(A)** GDF-15 expression in UACC 903 and B16F10 cells grown on collagen ECM or in suspension compared to adherent control conditions. Western blot analysis shows that both altered ECM context and anchorage independence induce GDF-15 expression. α-Tubulin was used as a loading control. (**B-1)** Schematic of the **“sandwich”** setup in which cells are first grown on glass coverslips, then the coverslip is inverted onto a glass–plastic support to position the cell layer between rigid surfaces. **(B-2)** Schematic of the **weight-based compression** assay in which defined masses are placed on the upper glass layer to apply graded normal forces and generate increasing compressive pressures on the cell monolayer. **(C)** Western blot analysis of GDF-15 expression in UACC 903 cells exposed to increasing mechanical pressure (22 Pa, 38 Pa, 54 Pa, 100 Pa), with the flipped coverslip alone (22 Pa) serving as a low-pressure condition and UACC 903M (red) as a positive control. **(D)** Quantification of fold change in GDF-15 expression relative to the uncompressed negative control in UACC 903 cells, demonstrating a pressure-dependent increase in GDF-15. **(E)** Western blot analysis of GDF-15 expression in B16F10 cells under the same graded mechanical pressures, with UACC 903M (red) used as a positive control. **(F)** Quantification of fold change in GDF-15 expression relative to the uncompressed negative control in B16F10 cells, showing a similar pressure-dependent induction of GDF-15.

### GDF-15 Synergizes with Vascular Mediators to Enhance Endothelial Permeability and Promote Metastasis

While GDF-15 has been implicated in immune modulation and metastatic cancer progression (12,17,18,27), its functional role in regulating tumor-associated vascular permeability remains undefined, and mechanisms enabling these processes are unknown. Given that melanoma often exhibits leaky (28), and disorganized vasculature (29), the role of GDF-15 in facilitating this process is unclear, and whether it would facilitate immune evasion and metastatic dissemination is unknown. Therefore, understanding how the tumor microenvironment, via GDF-15, drives vascular dysfunction is critical. Notably, GDF-15 is secreted in response to mechanical stress. It is elevated in advanced melanoma (30), yet it remains unknown whether it directly influences endothelial integrity or interacts with classical vascular permeability mediators, such as VEGF (31), bradykinin (BK) (32), or platelet-activating factor (PAF) (33) that are present in tumors.

To address this knowledge gap, we investigated whether GDF-15 acts in concert with inflammatory signals constitutively present in tumors (34) to disrupt endothelial junctions and promote vascular leakiness. The following experiments identify a previously unrecognized synergy between GDF-15 and these mediators, providing new insight into how the mechanical and inflammatory components of the TME converge to compromise vascular barriers and potentially promote metastatic spread.

Initial experiments used a confluent monolayer of human endothelial cells (HUVECs) (35) treated with GDF-15 alone or in combination with vascular permeability factors, BK or VEGF, typically present in tumors. This treatment significantly increased gap formation within the endothelial layer. Representative images highlight gaps (red arrows) in treated cells relative to controls (**Figures 4A, B**). Quantification confirmed that 1 µg/ml GDF-15, in combination with 7.8-125 nM BK or 12.5-100 ng/ml VEGF, synergistically enhanced gap formation, demonstrating a cooperative effect. These findings suggest that GDF-15 contributes to endothelial leakiness by interacting with vascular mediators present in tumors.

**Figure 4.**
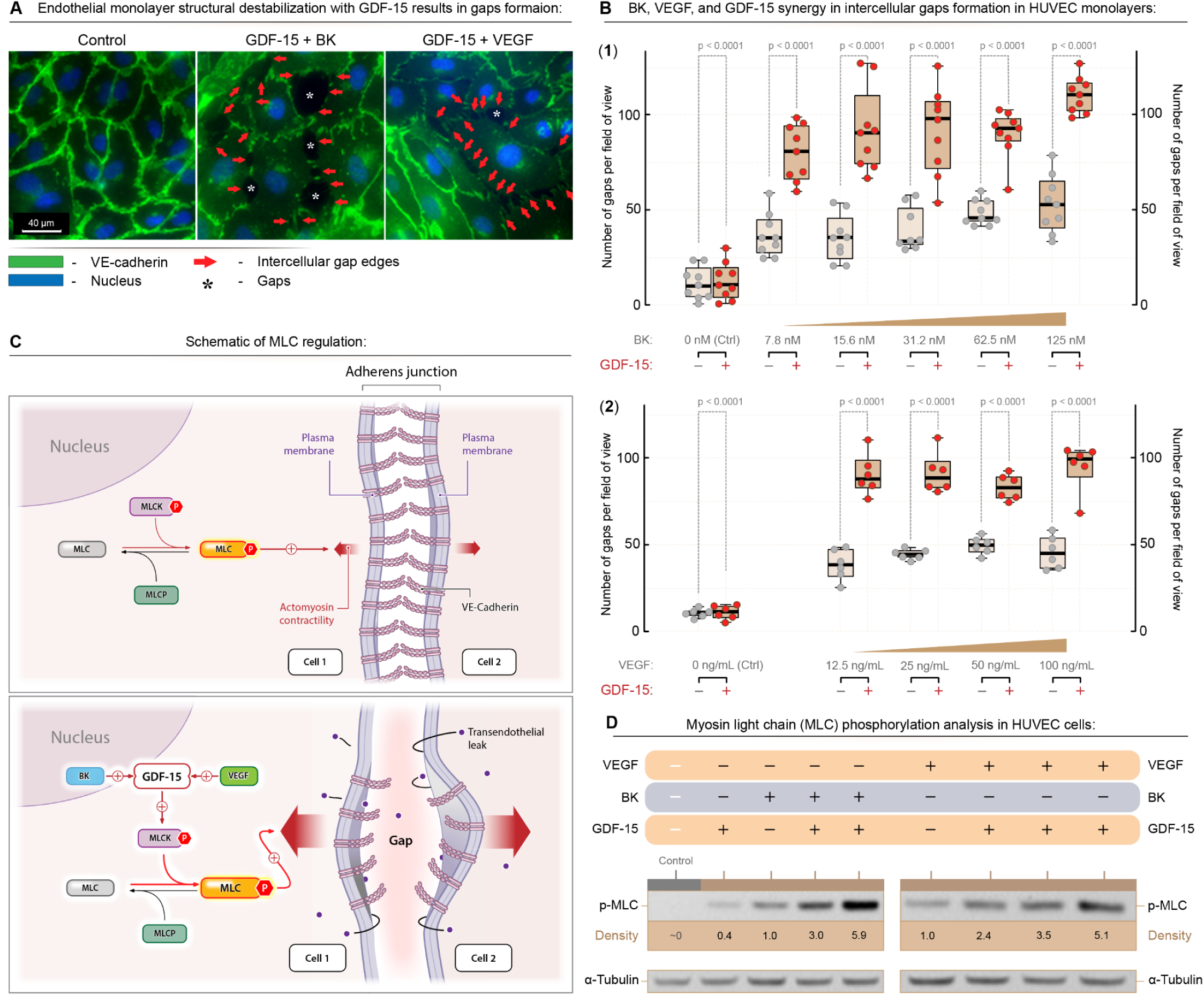
GDF-15 synergizes with vascular permeability mediators to enhance gap formation in an endothelial cell monolayer through increased pMLC-dependent cell contractility. **(A)** GDF-15 combined with either bradykinin (BK) or VEGF markedly increases intercellular gaps in confluent HUVEC monolayers compared to control, with red arrows indicating gap edges and asterisks marking representative gaps (40× magnification). **(B)** Quantification of intercellular gap numbers per field of view in HUVEC monolayers treated with increasing concentrations of BK (B1) or VEGF (B2) in the presence or absence of GDF-15, demonstrating a synergistic effect of GDF-15 with each permeability mediator on gap formation. **(C)** Schematic illustrating how GDF-15, together with BK or VEGF, enhances myosin light chain kinase (MLCK)-driven phosphorylation of myosin light chain (pMLC), thereby increasing actomyosin contractility and destabilizing VE-cadherin-based adherens junctions to promote gap formation. **(D)** Western blot analysis of HUVEC lysates showing that combined treatment with GDF-15 and BK or VEGF elevates pMLC levels relative to single treatments, consistent with enhanced contractility and increased endothelial gap formation. *In-vivo* permeability changes by GDF-15 and permeability mediators

To investigate the mechanism mediating GDF-15-mediated gap formation, endothelial cell contractility was examined with a focus on the phosphorylation of myosin light chain (MLC), whose signaling is reported to trigger cell contractility (35,36)(**Figure 4C**). Treatment with GDF-15, combined with inflammatory factors BK or VEGF, led to a dose-dependent increase in phospho-MLC levels, as shown by representative quantified Western blot images (**Figure 4D**), consistent with a mechanism in which GDF-15 synergizes with inflammatory factors to enhance cytoskeletal contraction in the endothelial cell layer. The contraction caused by phospho-MLC appeared to pull the edges of endothelial cells apart, forming gaps in the monolayer (35). These results highlight a novel function: GDF-15 cooperates with tumor-derived vascular permeability mediators to promote endothelial cell contraction and gap formation, implicating GDF-15 in vascular permeability.

To examine the effects of GDF-15 on vascular permeability in animal models, the Miles assay in mice was used; GDF-15 treatment, combined with BK or VEGF, significantly increased vascular permeability. This was evidenced by the enhanced retention of Evans blue dye leaking from the capillaries in the ears of treated mice, indicating increased vascular permeability (**Figure 5A**). The dye was retained in the ears when the endothelial gaps closed and could be objectively quantified. Quantitative analysis revealed a synergistic effect when GDF-15 was combined with PAF, compared with individual treatments (**Figure 5B**). Importantly, pharmacological intervention with ASR396, a sphingosine-1-phosphate receptor agonist reported to stabilize endothelial barriers (35), completely reversed the vascular leakiness induced by GDF-15 and PAF. This reversal strongly confirmed the specificity of GDF-15’s action in compromising vascular integrity.

**Figure 5.**
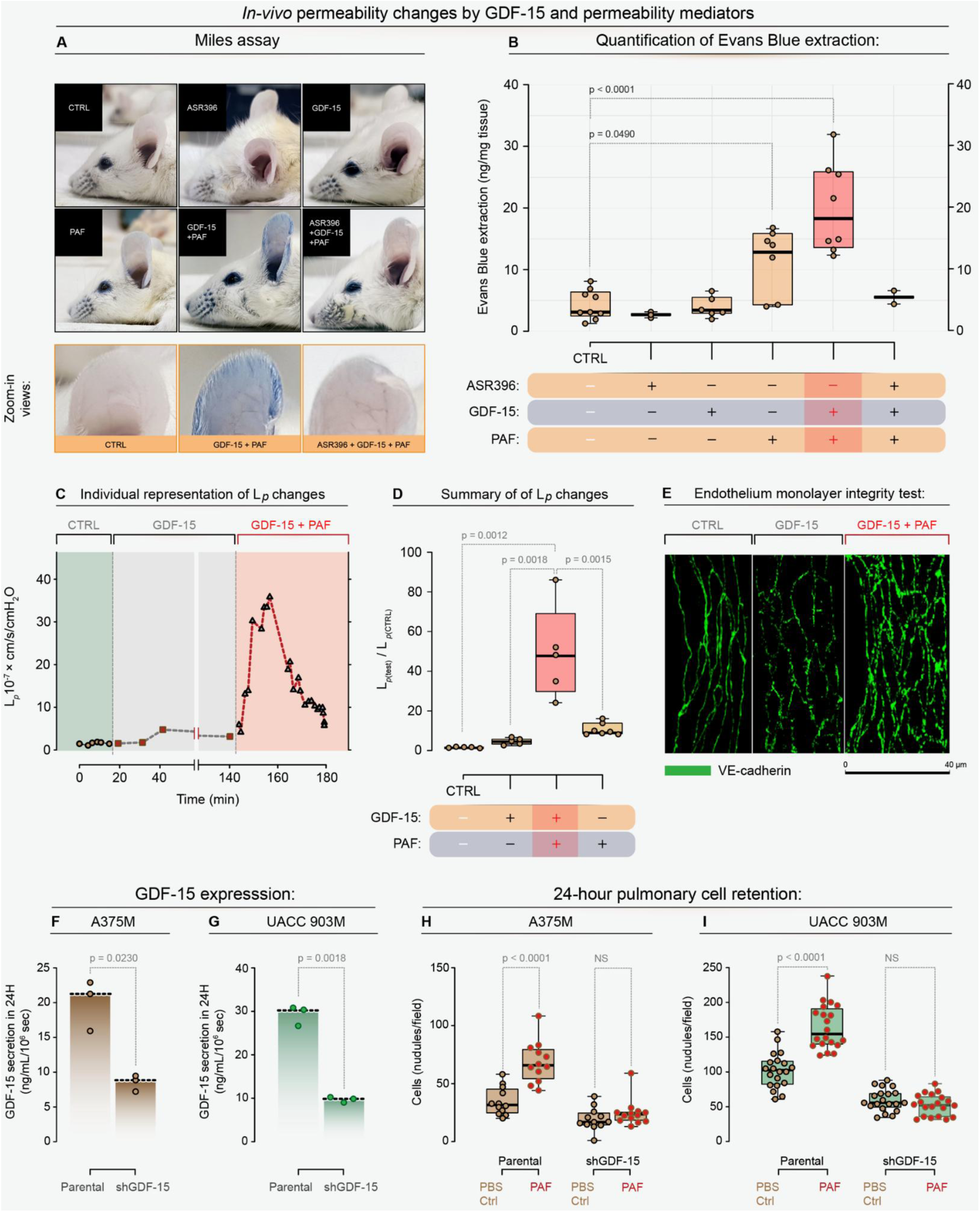
GDF-15 synergizes with vascular permeability mediators to enhance vascular leakiness in vivo, thereby promoting metastatic extravasation. **(A)** Representative Miles assay images showing Evans Blue dye accumulation in murine ears treated with GDF-15, PAF, or their combination, with GDF-15 plus PAF producing the most significant leakage, which is abrogated by co-treatment with the S1PR1 agonist ASR396. **(B)** Quantification of Evans Blue extracted from ear tissue, confirming that GDF-15 potentiates PAF-induced vascular leakage and that ASR396 reverses this effect. **(C)** Representative single-micro-vessel experiment in which rat mesenteric venules were sequentially perfused with control BSA Ringer’s solution, GDF-15 (10 µg/mL), and then GDF-15 plus PAF (10 nM), illustrating dynamic changes in hydraulic conductivity (*L_p_*) over time. **(D)** Summary of micro-vessel permeability measurements showing that GDF-15 alone does not significantly increase *L_p_*, but markedly augments the PAF-induced rise in *L_p_*, from baseline to approximately 50-fold over control (*L*_p test_ / *L*_p CTRL_). **(E)** Confocal VE-cadherin staining of fixed mesenteric micro-vessels revealing continuous junctions in control vessels, junctional disruptions after PAF, and extensive junctional breaks and gaps after combined GDF-15 plus PAF, consistent with the magnitude of permeability increase (scale bar, 40 µm). **(F,G)** ELISA of conditioned media demonstrating reduced GDF-15 secretion in A375M shGDF-15 and UACC 903M shGDF-15 cells compared with their respective parental lines. **(H,I)** Quantification of tumor cell retention in the lungs 24 hours after intravenous injection, showing that PAF enhances lung retention of parental A375M and UACC 903M cells but not GDF-15 knockdown cells, indicating that GDF-15 is required for PAF-driven metastatic extravasation.

For additional rigor, the effect of GDF-15 on vascular permeability was assessed in an individually perfused intact rat mesenteric venular microvessel model (35), by measuring one permeability coefficient, hydraulic conductivity (*L_p_*). **Figure 5C** shows the time course of the *L_p_* changes from an individual experiment when the vessel was perfused with control perfusate (Ringer’s solution with 1% BSA), GDF-15 (10 µg/ml), and GDF-15 plus PAF (10 nM). **Figure 5D** shows the data summary for the combination treatment, as well as the PAF perfusion alone-induced *L_p_* increases. Exposure of microvessels to GDF-15 alone did not cause significant increases in *L_p_*, but its exposure markedly potentiated the vascular responses to PAF, indicating a crucial priming role of GDF-15 in inflammatory mediator-induced permeability increases in intact microvessels. The PAF-induced peak increase with and without the pre-exposure of GDF-15 was 49.32 ± 10.52 *vs.* 11.10 ± 1.13 times that of the control, respectively. Images of VE-cadherin staining of rat microvessels showed a continuous, smooth distribution between endothelial cells under control conditions, frequent breaks at the peak increase in *L_p_* during PAF perfusion, and more pronounced gaps at the LP peak with perfusion of GDF-15 + PAF (**Figure 5E**). These images indicate that redistribution of adherens junction proteins occurs when microvessel permeability increases, and the extent of the breaks correlates with the magnitude of the permeability increase.

The impact of GDF-15 synergizing with vascular permeability mediators on metastasis, particularly extravasation, was investigated using an experimental lung metastasis model. Melanoma cells overexpressing GDF-15 (UACC 903M and A375M cells) were compared to cells with GDF-15 knockdown (UACC 903MshGDF-15 and A375MshGDF-15 cells), by quantifying the number of retained cells in the lungs 24 h after intravenous cell injection. GDF-15 ELISA assays confirmed a threefold reduction in secretion in knockdown cells (**Figures 5F, G**). Mice injected with GDF-15-expressing cells showed significantly higher extravasation into the lung parenchyma and subsequent retention of melanoma cells in the lungs compared to those injected with knockdown cells (**Figures 5H, I**). Notably, the presence of PAF further enhanced cell retention in the lungs, demonstrating that GDF-15 synergizes with PAF to create vascular gaps that facilitate tumor cell migration and metastasis. These results underscore the crucial role of GDF-15 in promoting metastatic conduits through its interactions with vascular permeability factors.

### Effect of GDF-15 on Tumor Development, Structure, and Suppression of NK Cell-Mediated Immunity

Despite increasing recognition of GDF-15 as a stress-responsive cytokine (37), ), its precise functional role in shaping the tumor microenvironment, particularly in promoting tumor growth, vascular disorganization, and immune evasion, remains unknown. While previous studies have identified GDF-15 serum levels as a potential biomarker for advanced melanoma (11), it has not been mechanistically linked to the hemorrhagic, highly vascularized phenotype observed in aggressive melanomas, nor to suppression of innate immune responses. We directly address these knowledge gaps by examining the effects of GDF-15 in driving a disordered, leaky vascular phenotype characterized by increased hemorrhagic areas, likely through its synergy with vascular permeability mediators. Furthermore, we investigate whether GDF-15 suppresses NK cell–mediated cytotoxicity, thereby providing a dual mechanism: both a mediator of vascular dysfunction and a suppressor of innate immunity.

The role of GDF-15 in melanoma tumor morphology development was examined using UACC 903M and A375M melanoma cell lines and GDF-15 knockdown counterparts (UACC 903MshGDF-15 and A375MshGDF-15 cells). Tumors derived from cells with GDF-15 knockdown showed more than a 60% reduction in tumor development compared to GDF-15-expressing tumors in nude mice (**Figures 6A, B**). Visual and histological analysis revealed that tumors expressing GDF-15 were visibly bloodier, with a higher density of red blood cells, compared with tumors with GDF-15 knockdown (**Figures 6C, D, & E**). Quantification of hemorrhagic areas in H&E-stained tumor sections showed a 50% reduction in UACC 903M knockdown tumors and over an 80% reduction in A375M knockdown tumors, highlighting the impact of GDF-15 on tumor hemorrhagic vascular morphology (**Figure 6E**). Thus, knockdown of GDF-15 resulted in smaller, less hemorrhagically vascularized tumors with reduced blood volume, suggesting that GDF-15 contributes to vascular disorganization associated with hemorrhagic areas.

**Figure 6.**
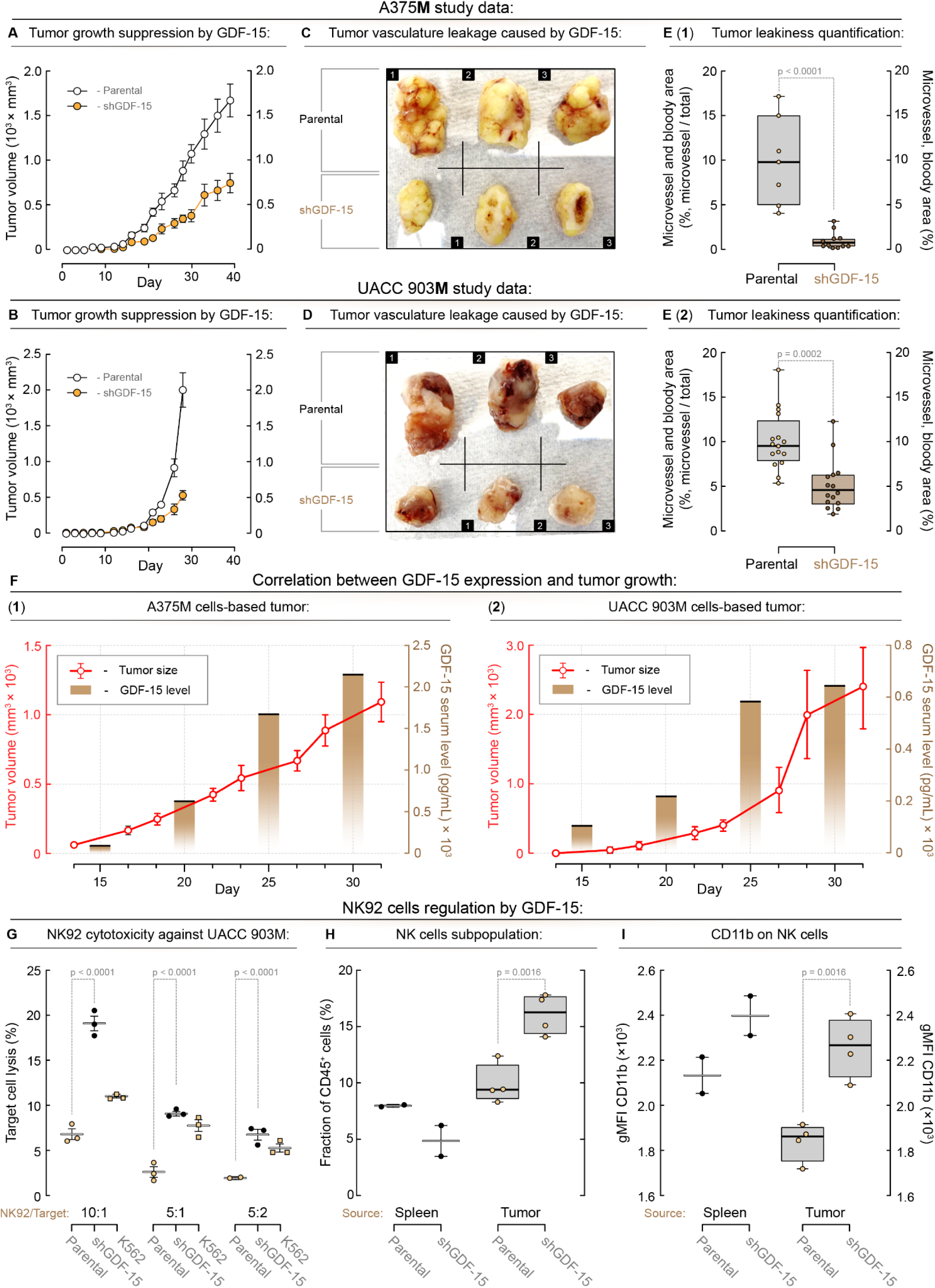
GDF-15 enhances tumor size by promoting vascular leakiness through synergism with endogenous vascular permeability mediators that regulate tumor hemorrhagic vascular morphology, thereby increasing tumor blood volume. **(A)** A375M and **(B)** UACC 903M parental cells expressing GDF-15 formed 60-70% larger tumors compared to shGDF-15 counterparts. Three representatives of **(C)** A375 M and **(D)** UACC 903M parental cells, compared with shGDF-15 tumors, showed gross differences in size and bloodiness. **(E)** Tumors shown in **(C** and **D)** were processed for H&E stain, and the areas of tumor hemorrhagic vascular morphology were quantified, showing significantly less in the parental tumors compared to the GDF-15 knockdown tumors. **(F)** Levels of GDF-15 in serum measured by ELISA increased with the size of A375M and UACC 903M tumors. **(G)** GDF-15 decreased NK cell-mediated killing of tumor cells. Target UACC 903M parental or shGDF-15 cells were labeled with 5 µM Cell Trace Violet (CTV) and then combined with NK-92 cells at the indicated effector to target ratios (E:T) in triplicate for 5 h. Cultures were stained with 7AAD for 10 min at room temperature, and the percentage of live target cells (7AAD-negative, CTV+) in each culture was quantified by flow cytometry. UACC 903M parental and shGDF-15 tumors from nude mice were subjected to NK cell profiling. **(H)** Significantly higher percentages of NK cells were present in the shGDF-15 tumors. **(I)** Higher NK cell activity was also observed in the shGDF-15 tumors. CD11b was used as an NK activation marker.

To demonstrate that tumor burden correlates with serum GDF-15 levels, an ELISA was performed on serum samples from mice bearing UACC 903M and A375M tumors of increasing size. Results showed a direct relationship between tumor size and serum GDF-15 levels (**Figure 6F**). In A375M tumors, GDF-15 levels increased from approximately 200 pg/mL to 2,000 pg/mL as tumor volume increased from 200 mm³ to 500 mm³. Similarly, UACC 903M tumors exhibited serum GDF-15 levels of approximately 100 pg/mL at 300 mm³, rising to approximately 1,500 pg/mL at 2,000 mm³. These data demonstrate that GDF-15 is secreted in proportion to tumor growth and provide strong evidence that GDF-15 enhances tumor hemorrhagic vascularization and metastatic progression. The bloody and disorganized vascular phenotype in tumors expressing GDF-15 supports its role in creating a permissive tumor microenvironment. Combined with its correlation with tumor size and elevated serum levels in melanoma patients, GDF-15 emerges as a critical mediator of melanoma progression and a potential biomarker and therapeutic target.

The role of melanoma cell GDF-15 in suppressing immunity was assessed using an in vitro co-culture model to perform a natural killer cell (NK92) cytotoxicity assay. Human NK92 NK cells were co-cultured with cell trace violet-labeled UACC 903M cells, either constitutively expressing or deficient in GDF-15, at defined effector-to-target ratios, and the efficiency of target cell killing was determined (**Figure 6G**). The NK-sensitive target cell K562 was used to validate the dose-dependent activity of the NK92 cells. Knockdown of GDF-15 significantly increased NK92-mediated cytotoxicity by approximately threefold (**Figure 6G**).

The immune suppression effect of GDF-15 was further confirmed in UACC 903M and UACC 903MshGDF-15 tumors. Thirty-six days after cell injection, UACC 903M and UACC 903MshGDF-15 tumors from nude mice were subjected to NK and myeloid cell profiling. There was a significant increase in NK cell infiltration and activity in UACC 903MshGDF-15 tumors (**Figure 6H & I**). Also, a substantial increase in macrophages was observed (**Supplementary Figure S3**). These findings provide strong evidence that GDF-15 contributes to immune evasion by impairing NK cell cytotoxicity. Together, the in vivo and in vitro data underscore the pivotal role of GDF-15 in creating an immunosuppressive environment that supports tumor progression. Targeting GDF-15 may offer a therapeutic strategy to restore NK cell function and enhance anti-tumor immunity in melanoma.

## DISCUSSION

Our study identifies GDF-15 as a mechanically induced mediator of melanoma progression, revealing new dimensions to its role in shaping tumor structure, promoting invasion, and suppressing immune surveillance. While previous reports have established that GDF-15 exerts pleiotropic effects, including promoting epithelial-to-mesenchymal transition (EMT) (12), regulating angiogenesis (38), and modulating the ECM(39), the mechanisms by which GDF-15 contributes to melanoma-specific immune evasion and tumor architecture remain poorly understood. Our findings significantly extend these observations by showing that GDF-15 expression is regulated by TME rigidity, as recapitulated in GelMA microgels that partially mimic the ECM. Furthermore, the forces exerted on cells during tumor progression would also increase GDF-15 levels, contributing to changes in tumor architecture and immune suppression, which could be especially relevant in the context of acral melanoma (40), which develops in high-mechanically stressed, rigid collagen regions of the body (40).

Our findings demonstrate that GDF-15 is highly expressed in melanoma via mechanical induction mediated by ECM rigidity and mechanical forces that shape tumor development, and that this expression correlates with significant alterations in tumor morphology and vascularization. Tumors overexpressing GDF-15 exhibited disorganized hemorrhagic vascular networks leading to increased blood volume, suggesting that GDF-15 plays a critical role in hemorrhagic tumor vascularization (14,41–43). This observation is consistent with previous studies linking GDF-15 to aberrant angiogenesis (14), but our mechanistic analysis of the disorganized tumor vasculature provides deeper insight into this relationship. The reduction in the vessel density, leakage, and chaotic architecture following GDF-15 knockdown highlights the therapeutic potential of targeting GDF-15 to restore vascular integrity and reduce tumor progression.

A significant and novel finding of this study is the identification of a synergistic interplay between GDF-15 and other vascular permeability factors within the tumor microenvironment, highlighting a previously unrecognized mechanism driving tumor progression. Our results demonstrate that GDF-15 does not act in isolation but rather cooperates with other pro-angiogenic and vascular-disrupting factors to enhance vascular permeability and disorganization, ultimately promoting tumor growth and metastasis (44). This synergism was particularly evident in the pronounced vascular abnormalities observed in GDF-15-expressing tumors, including increased vessel density, leakage, and chaotic architecture, as compared to tumors with GDF-15 knockdown. These findings suggest that GDF-15 amplifies the effects of established vascular permeability factors, such as VEGF (14), to create a permissive environment for tumor expansion and immune evasion. By linking GDF-15 to these processes, this study provides the first evidence of its cooperative role in vascular remodeling, positioning GDF-15 as a critical integrator of vasculogenic signals within the tumor milieu.

This discovery opens new avenues for therapeutic intervention by targeting GDF-15 in conjunction with established anti-angiogenic therapies (45), offering the potential to restore vascular integrity and improve drug delivery while simultaneously mitigating immune suppression. By demonstrating that GDF-15 knockdown not only reduces hemorrhagic tumor vascularization but also improves immune function, this research highlights the cytokine as a versatile therapeutic target. The therapeutic implications extend beyond melanoma, as GDF-15 is implicated in several cancers and pathological conditions involving immune suppression and aberrant angiogenesis(8,14,41–43).

The ability of GDF-15 to suppress immune responses (8), particularly those mediated by natural killer (NK) cells, underscores its role in creating an immunosuppressive TME. particularly those mediated by natural killer (NK) cells, underscores its role in creating an immunosuppressive TME. This is supported by cytotoxicity assays showing diminished NK92-mediated killing of melanoma cells expressing GDF-15, reinforcing the cytokine’s immunosuppressive properties. This suggests that GDF-15 acts as a key mediator of immune evasion, specifically impairing NK cell-mediated cytotoxicity. These findings add to the growing body of evidence implicating GDF-15 in modulating anti-tumor immunity (17,18), providing strong justification for its therapeutic targeting to enhance immune surveillance. Mechanistically, GDF-15 has been shown in other cancer types to impair T cell recruitment by blocking LFA-1-dependent adhesion, thereby limiting T cell extravasation and infiltration into tumors (17). This immune exclusion is a major contributor to resistance against immune checkpoint inhibitors (ICIs), such as anti-PD-1 therapies (17). Beyond T cells, GDF-15 has also been reported to modulate the myeloid compartment in other cancers by inhibiting dendritic cell maturation, suppressing M1 macrophage polarization, and promoting a tolerogenic, anti-inflammatory phenotype in the TME (8). This broad immunosuppressive effect further impairs anti-tumor immunity and supports tumor progression.

Numerous studies have reported that serum levels of GDF-15 were significantly elevated in patients of multiple cancer types, including melanoma, compared to healthy individuals (46–48). In this report, we show that GDF-15 levels increased in proportion to tumor burden. This correlation highlights the potential utility of serum GDF-15 as a non-invasive biomarker for tumor burden, disease progression, and therapeutic response monitoring (11,30). Furthermore, the differential expression of GDF-15 in melanoma cell lines derived from distinct disease stages underscores its role in driving tumor progression and metastasis (14), suggesting that GDF-15 may serve as a key factor in the transition to more aggressive melanoma phenotypes. Elevated serum and tumor GDF-15 levels are predictive of poor response to ICIs and shorter overall survival in melanoma and other cancers (11,49). Neutralization of GDF-15 with antibodies (e.g., visugromab) restores T-cell infiltration, enhances anti-tumor immune responses, and improves the efficacy of ICIs in preclinical models and early-phase clinical trials (50). These findings have led to ongoing clinical trials evaluating GDF-15 blockade in combination with ICIs in patients with refractory or resistant tumors (50). GDF-15 also promotes tumor cell proliferation, migration, invasion, and EMT via pathways such as PTEN/PI3K/AKT, thereby contributing to tumor aggressiveness and therapy resistance (30). Its role as a biomarker is supported by consistent associations between high GDF-15 levels and advanced disease stage, metastasis, and immune exclusion (11,30,46–48). Additionally, GDF-15 is implicated in cancer cachexia, and its inhibition may improve both immune response and patient quality of life (19,51–53).

From a translational perspective, the clinical significance of these discoveries is that GDF-15 may serve not only as a biomarker for tumors with high mechanical stress and immune exclusion but also as a therapeutic target to reprogram the TME toward a more immuno responsive state. Given the dismal outcomes in patients with immune-resistant acral melanoma and the limited efficacy of immune checkpoint inhibitors in this setting, targeting GDF-15 could provide a new avenue to enhance the effectiveness of current immunotherapies (50). Furthermore, GDF-15 expression may stratify patients who are less likely to respond to checkpoint inhibitors alone (49) and might benefit from combination strategies incorporating GDF-15 blockade (50,53).

In conclusion, this study identifies GDF-15 as a mechano sensing-modulated regulator of tumor architecture and immune suppression, uncovering a previously unrecognized link between mechanical rigidity and external force, synergy between GDF-15 and mediators of vascular permeability, disorganized vascular hemorrhagic remodeling, and immune evasion, thereby revealing a therapeutic vulnerability across both common and rare melanoma subtypes. By illuminating new functions and processes regulated by GDF-15, these findings fill critical knowledge gaps in the field and open the door to novel therapeutic approaches designed to overcome immune resistance and improve outcomes for patients with common and rare melanoma types.

## MATERIALS AND METHODS

### Cell lines and culture conditions

The human cutaneous melanoma cell lines, A375P, A375M/A375M-GFP; constitutively expressing GDF-15, UACC 903, UACC 903M/UACC 903M-GFP; constitutively expressing GDF-15, and WM4235 (acral melanoma) and mouse melanoma cell lines, B16-F10 and M4 (a kind gift from Dr. Glenn Merlino, NIH)(26) cells were maintained in DMEM (Invitrogen) supplemented with 5% FBS (Atlantic Biologicals) and GlutaMAX (Life Technologies). GDF-15 expression was knocked down in UACC 903M and A375M cells by transfecting cells with a lentiviral vector containing GDF-15 shRNA PLASMIDS (TRCN0000058392) and selecting for 7 days with puromycin. Human umbilical vein endothelial cells (HUVEC)(Lonza) were maintained on gelatin-coated plates in endothelial growth media (EGM) supplemented with 2% FBS (Lonza). Melanoma cells were used within 20 passages; HUVEC cells were used between passages 2-5. Bradykinin (BK) and PAF were from Sigma, while GDF-15 and VEGF were from Peprotech.

### NK Cell Cytotoxicity Assay

Target melanoma cells were resuspended at 10^6^/ml in PBS(Cytiva) containing a 1:1000 dilution of CellTrace™ Violet (Molecular Probes) and incubated for 20 min at 37°C. The labeling reaction was quenched by adding 10 ml of complete RPMI-1640 containing 10% FBS, then centrifuged and resuspended to 2×10^5^/ml in complete RPMI-1640/10% FBS. Target cells were plated at 2×10^4^/well in 96-well round-bottom plates. NK-92 cells (ATCC CRL-2407) were grown in RPMI-1640 containing 10% FBS and 15% Horse Serum (Gibco) with 10 ug/ml recombinant human IL-2 (Peprotech). NK-92 cells were passaged 1:5 every 3 days. On the day of the assay, NK-92 cells were harvested, resuspended at 10^6^/ml in RPMI-1640/10 % FBS, and serially diluted 2-fold before adding 100 µL/well in triplicate to wells containing target cells for a final effector: target ratio of 10:1, 5:1 and 2.5:1. Control wells contained only target cells and an equivalent amount of media to determine the baseline frequency of dead cells. Cultures were incubated at 37°C, 5% CO_2_ for 4 h, after which 7-AAD (Invitrogen) was added at 2.5 µL/well for 10 min at room temperature. Samples were run on a BDFACSymphony A3 in the Penn State Flow Cytometry Core and data analyzed using FlowJo software (v.10.10). Percentage dead target cells (7-AAD+ of cell trace violet+) were determined by subtracting the mean value of target cells alone from each value acquired from wells containing both targets and NK-92 cells.

### GDF-15 sandwich enzyme-linked immunosorbent assay (ELISA)

Cells were cultured to confluency, washed with PBS, and fresh media was added. 24 h later, conditioned media were collected from the plates and analyzed by GDF-15 ELISA for quantification of protein levels in cell culture media and serum samples, using the DOST ELISA for human GDF-15 (R&D Systems) according to the manufacturer’s protocol. Cells were counted at the end of the experiment to normalize GDF-15 expression. Assays were replicated at least twice.

### GelMA synthesis, GelMA droplet fabrication, formation of individually photocrosslinked microgels, and microgel stability and rheological characterization are described in detail in the Supplementary Information

Procedures followed those listed in our previous publications (54,55) or cited in references listed in the supplementary information section.

### Endothelial cell monolayer gap formation assay

HUVECs were seeded onto coverslips coated with fibronectin (Sigma) and cultured to confluence. Media were changed to EGM with 0.5% FBS for at least 18 h prior to treatment. GDF-15 was added to HUVECs at 1 mg/ml for 1 h, after which the cells were stimulated with 0.5 µM bradykinin (BK) for 30 min and with 100 ng/ml vascular endothelial growth factor (VEGF) for 1 h. HUVECs were fixed in 2% cold paraformaldehyde (Electron Microscopy Sciences) at room temperature for 30 min, then blocked in 1% BSA (Rockland Immunochemicals) at room temperature for 30 min, and permeabilized with 0.1% Triton X-100 (Fisher) for 5 min. Goat anti-human VE-cadherin (Santa Cruz) diluted in 1:100 was incubated with cells at 4°C overnight. Alexa488 donkey anti-goat antibody (Invitrogen) diluted in 1:300 was incubated with cells at room temperature for 1 h in the dark. Coverslips were mounted on slides using the antifade mounting medium with DAPI (Vector Laboratories) and stored at 4°C in the dark before imaging by fluorescence microscopy.

### Western blot analysis

GDF-15 was added to HUVECs 1 h before 0.5 mM BK and 100 ng/ml VEGF were added for 5 and 15 min, respectively. Cells were lysed in RIPA buffer containing protease and phosphatase inhibitors (Pierce Biotechnologies). 20 µg of cell lysate was used for western blotting and probed with antibodies according to the suppliers’ recommendations. Antibodies to phosphor-myosin light chain (pMLC) (Cell Signaling) and to alpha-tubulin (Sigma) for use as a loading control were used for western blotting.

For western blot analysis of melanoma cells cocultured with microgels. Melanoma cells were co-cultured with microgels at a 1:3 cell-to-microgel ratio. After 24 h, cells were collected and centrifuged, and the resulting pellet was incubated with collagenase (1 mg/mL) at 37°C for 30 min to digest the microgels. Following digestion, the pellet was resuspended in RIPA buffer containing protease and phosphatase inhibitors (Pierce Biotechnologies). Then, 25 µg of cell lysate was probed with antibodies according to the suppliers’ recommendations. Antibodies for GDF-15 (D2A3) (Cell Signaling) and ERK2, used as a loading control (Santa Cruz).

### Immunofluorescence analysis of GDF-15 expression in melanoma cells co-cultured with microgels

Melanoma cells were co-cultured with both stiff and soft microgels at a 1:3 cell-to-microgel ratio for 2 h. Following a 24-h incubation, cells were fixed in a mixture of 3% paraformaldehyde, 0.25% Triton X-100, and 0.2% glutaraldehyde. The fixation mixture was prepared by combining 750 µL of 4% paraformaldehyde (PFA), 2.5 µL of Triton X-100, 8 µL of 25% glutaraldehyde, and 240 µL of 1× PBS (without calcium/magnesium). Cells were incubated in this solution for 15 min at 37°C, followed by two 10-minute washes with 1×PBS. To reduce autofluorescence, cells were quenched using a cold cytoskeleton buffer supplemented with freshly added sodium borohydride (1 mg/mL) for 15 min on ice, followed by three additional PBS washes. Immunostaining was performed using a primary GDF-15 antibody at a 1:100 dilution. Following primary antibody incubation, cells were treated with an Alexa Fluor 598-conjugated anti-rabbit secondary antibody. Hoechst dye was used for nuclear staining, and phalloidin was applied to visualize F-actin. Fluorescence imaging was performed using a confocal microscope with a 10× objective to assess GDF-15 expression under varying microgel stiffness conditions.

### ECM Stiffness and Anchorage Independence Assay

Melanoma cells were cultured under three different conditions to assess the impact of ECM stiffness and anchorage independence on GDF-15 expression. For the ECM condition, cells were seeded on Cytosoft Discovery substrates (Advanced Biomatrix) with a defined stiffness of 2 kPa collagen (soft matrix). For the suspension condition, cells were plated on PolyHEMA-coated dishes to prevent attachment, mimicking anchorage-independent growth. Adherent cells grown under standard tissue culture conditions served as the control. All cultures were maintained for 24 h, after which cell lysates were collected using standard lysis protocols and processed for Western blot analysis to assess GDF-15 expression.

### Weight-Based Compression Force Assay

Weight-Based Compression Force Assay. Melanoma cells were seeded onto sterile 25 mm × 25 mm No. 1 glass coverslips placed in a culture plate and allowed to adhere for 24 h under standard culture conditions. After 24 h, the coverslips were gently inverted so that the cell-coated surface faced downward onto the well bottom, thereby creating a confined space in which mechanical compression could be applied by placing defined masses on top of the coverslip. To quantify the resulting compressive stress, the vertical force generated by the coverslip and any additional weights was calculated as the product of their masses and gravitational acceleration.

For the coverslip alone (mass ≈ 1.4 g), the baseline force was taken as the planar area of the coverslip, and the corresponding baseline compressive pressure experienced by the cells was obtained from

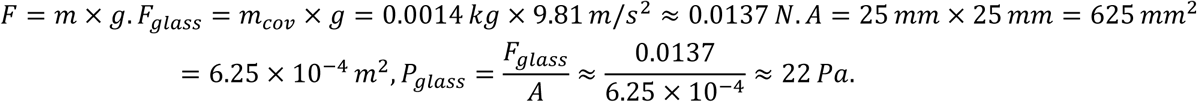

This value represents the constant gravitational load of the inverted coverslip even in the absence of externally added weights (flipped control condition).

For experiments with additional mechanical loading, calibrated masses of 1 g, 2 g, or 5 g were placed centrally on the inverted coverslip to increase the compressive force on the cell monolayer. The incremental force for each condition was calculated using the same relation *F* = *m* × *g*; for example,

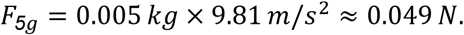

The total force acting on the cells was then the sum of the baseline coverslip force and the weight-derived force. The corresponding total pressure was (For the 5 g condition).

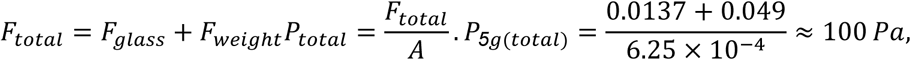

The pressures for the 1 g and 2 g conditions were calculated analogously by substituting the appropriate masses. In all cases, forces were assumed to be uniformly distributed across the coverslip area. A flipped control (inverted coverslip with no added weight, experiencing only *P*_*glass*_) and an unflipped/negative control (standard culture, no compression) were included in parallel. After 24 h of exposure to the indicated compressive conditions, cells were lysed using standard buffer and harvested for Western blot analysis of GDF-15 expression.

### Animal Studies

The Pennsylvania State University IACUC approved all procedures undertaken in animals. Both male and female of mice (8-10 weeks old) was used in studies unless specified otherwise.

### Melanoma cell lung retention assay

100,000 GFP-tagged human melanoma cells, UACC 903M, UACC 903M shGDF-15, A375M, and A375M shGDF-15, were intravenously injected into NRG mice 30 minutes before mice were intraperitoneally injected with 0.5 mg PAF. Mice were sacrificed after 24 h, and lungs were removed and imaged by fluorescent microscopy to quantify the number of retained cells.

### Miles vascular permeabilit assay

1 mg of GDF-15 was intraperitoneally injected for 2 h and/or sphingosine-1-phosphate receptor 1 agonist, ASR396, was intraperitoneally injected at 5 mg/kg for 30 mins, followed by 1 mg GDF-15 intraperitoneal injection for 2 h. Mice were then intravenously injected with 10 ng PAF diluted in 100 ml 1% Evans Blue and were sacrificed 20 mins later. Ears were collected and weighed, and 250 ml of formamide was added to each ear sample, which was incubated overnight at 55 °C. The peak Evans Blue absorbance at 620 nm was then measured.

### Measurement of hydraulic conductivity (*L_p_*) in individually perfused rat mesenteric micro-vessels and VE cadherin micro-vessel staining are described in detail in the Supplementary Information

A modified Landis technique was used to measure *L_p_* in individually perfused micro-vessels in the rat mesentery, as previously described (23).

### Microgel-based tumor modeling

UACC 903 cells and microgels were incubated at a 1:1 ratio for 24 h before injection into NRG mice. Tumor progression was monitored for seven days, after which tumors were extracted and processed for immunofluorescence analysis of GDF-15 expression. Extracted tumors were fixed in a fixation/permeabilization buffer containing 3% paraformaldehyde, 0.25% Triton X-100, and 0.2% glutaraldehyde. The fixation solution was prepared by combining 750 µL of 4% paraformaldehyde (PFA), 2.5 µL of Triton X-100, 8 µL of 25% glutaraldehyde, and 240 µL of 1× PBS (without calcium and magnesium). Tumors were incubated in this solution for 15 min at 37°C, followed by two 10-minute washes with 1× PBS. To minimize autofluorescence, samples were quenched using a cold cytoskeleton buffer supplemented with freshly added sodium borohydride (1 mg/mL) for 15 min on ice, followed by three additional PBS washes. Tumor sections were immunostained for GDF-15 expression with a primary antibody at a 1:100 dilution. Following primary antibody incubation, sections were treated with an Alexa Fluor 598-conjugated anti-rabbit secondary antibody. Hoechst dye was used for nuclear staining, and phalloidin was applied to visualize F-actin. Fluorescence imaging was performed using a confocal microscope with a 10× objective to assess GDF-15 expression under different microgel stiffness conditions.

### Tumor vascular studies

1 million UACC 903M, UACC 903M shGDF-15, A375M, and A375M shGDF-15 cells were subcutaneously injected into the flank of nude mice. Mice were monitored, and tumors were measured using a digital caliper twice a week. Mice were sacrificed when tumors from UACC 903M and A375M cells reached 1500 −2000 mm^3^ and tumors were removed and fixed with 10% formalin. Tumor sections were subjected to an H&E stain. Hemorrhagic vascular area distribution was quantified by microscopy.

### GDF-15 assessment with increasing tumor burden

UACC 903M and A375M cells were subcutaneously injected into female nude mice. Tumors were measured by a digital caliper twice a week. On days 15, 20, 25, and 30, 200 µl of blood was collected from the submandibular vein, and serum was used to measure GDF-15 levels by ELISA, as detailed above.

### Statistical analysis

All subjects were randomly assigned to experimental treatment groups. Data collection and subsequent quantification were performed in a double-blinded manner. Animals were monitored daily for signs of discomfort and distress. Mice were humanely euthanized and excluded from data collection if they lost 10% of their initial body weight or showed signs of lethargy or respiratory distress. Tumor data are shown as mean ± SEM. All data for GDF15 evaluation are presented as median ± SEM. All other data are shown as mean ± SD. A two-sided t-test was used to compare two groups; for experiments with more than two groups, one-way or two-way ANOVA was performed, followed by Dunnett’s, Tukey, or Bonferroni correction as appropriate. All experiments were repeated at least twice for increased rigor and robustness. A two-sided p<0.05 was considered significant.

## ACKNOWLEDGEMENT

This research was funded by NIH NCI grant R01CA241148-03 (G.R); NIH grant supplement RO1CA241148-0252-Ext (G.R); the Foreman Foundation for Melanoma Research (G.R); the Geltrude Foundation (G.R), and the Chocolate Tour Cancer Research Fund (G.R). HL144620 (P.H.) and NIH NINDS grant R01NS139128 (P.H.). Meghan Rose Bradley Foundation (A.S.), the Dorothy Foehr Huck and J. Lloyd Huck Early Career Chair (A.S.), and by NIH NHLB R01HL167939 (A.S.). We acknowledge the support and assistance of Lynne Beidler, Teah Batdorf, Virginia Robertson, Nazir A. Lone, Arati Sharma, Arun Sharma, Saba Keshavarz, and Krishne Gowda.

