## Supplementary Information for "Convergent Roles of Growth Differentiation Factor-15 (GDF-15) in Mechanotransduction, Vascular Disorganization, and Immune Suppression in Melanoma"

**Supplementary Figure Legends**

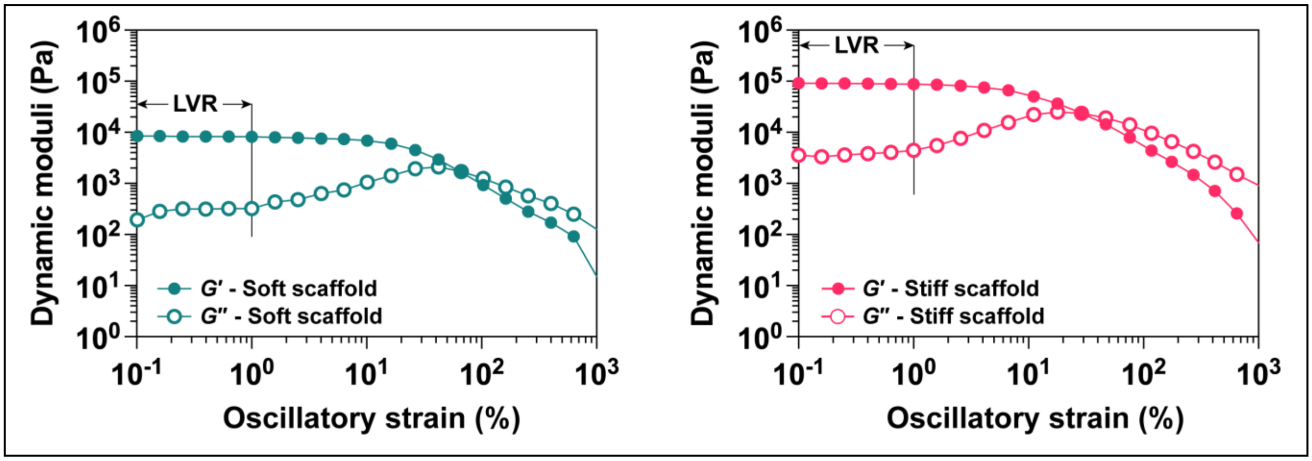


**Figure S1.** Oscillatory amplitude sweep tests conducted at a constant frequency (1 rad s-1) to assess the LVR of soft and stiff bulk hydrogel scaffolds, prepared with 4 or 15% w/v GelMA, respectively.

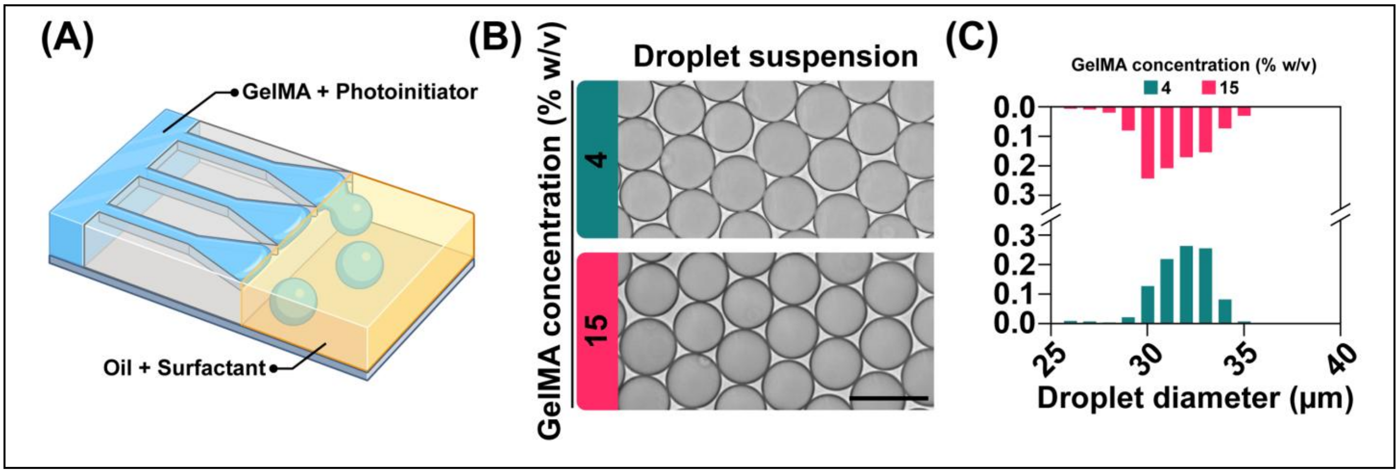


**Figure S2. Microgel droplet formation and stability analysis. (A)** A step-emulsification device is used to fabricate GelMA droplets. The dispersed phase consists of GelMA (4 or 15% w/v) and photoinitiator (0.1% w/v) in DPBS, while the continuous phase is composed of surfactant (2% v/v) in oil (Novec™ Engineered Fluid 7500). **(B)** Brightfield microscopy images of soft and stiff GelMA droplets, prepared with 4 and 15% w/v GelMA, respectively. Scale bar is 50 μm. **(C)** Size distribution of soft and stiff droplets formed using the microfluidic device (*n* > 1000).

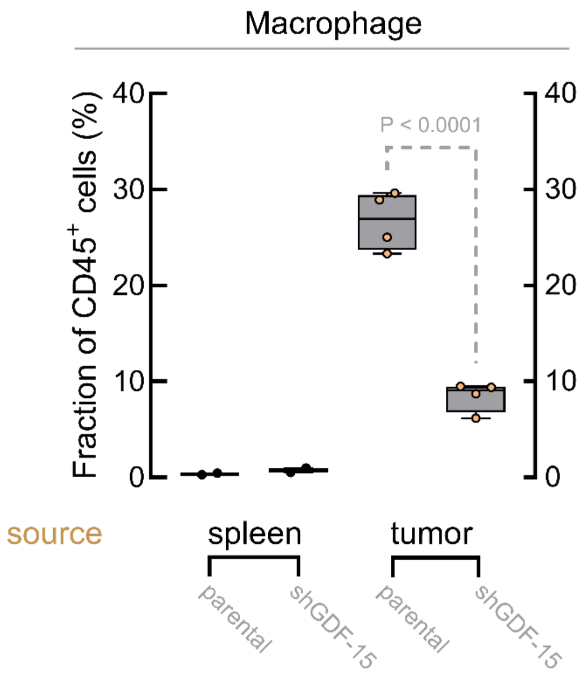


**Figure S3. Macrophage distributions in UACC 903M and UACC 903MshGDF-15 tumors.** UACC 903M and UACC 903MshGDF-15 cells were injected subcutaneously into nude mice. Tumors and spleens were harvested after 36 days. There was a significant increase in macrophages within UACC 903MshGDF-15 tumors.

**Supplementary Methods**

**GelMA synthesis, GelMA droplet fabrication, formation of individually photocrosslinked microgels and microgel stability.** Briefly, 100 mL of DPBS (MilliporeSigma) was heated to 50°C under continuous stirring. Then, 10 g of gelatin (Type A, ~300 g Bloom, from porcine skin, MilliporeSigma) was slowly added to the solution until fully dissolved. Next, 1.25 mL of methacrylate anhydride (MilliporeSigma) was added dropwise to the 10% w/v gelatin solution. The reaction was carried out for 2 h under stirring at 50°C and was halted by adding 200 mL of 50°C DPBS, while being stirred for an additional 15 min at 50°C. The solution was then poured into dialysis membrane tubing (12–14 kDa molecular weight cut-off, Spectrum Laboratories) and dialyzed against ultra-pure Milli-Q water (electrical resistivity ~18.2 MΩ cm at 25 °C, Millipore Corporation) at 40°C under stirring. After 7 days, 200 mL of 40°C Milli-Q water was added to the GelMA solution, while being stirred for 20 min, followed by sterile-filtration using a 0.2 μm vacuum filter (VWR). The filtered solution was then transferred to a -80°C freezer and kept for at least 48 h. The frozen GelMA solution was transferred to a Labconco FreeZone 4.5L Benchtop freeze dryer (Labconco) and lyophilized at vacuum pressure ~ 0.016 mbar until a solid GelMA was obtained.

Lyophilized GelMA was dissolved in 0.1% w/v lithium phenyl-2,4,6-trimethylbenzoylphosphinwere (LAP, MilliporeSigma) solution (40°C) to prepare 4 or 15% w/v solutions for the dispersed phase. For the continuous phase, a 2% v/v Pico-Surf® (Sphere Fluidics) solution was prepared in Novec™ 7500 Engineered Fluid (3M).

A step emulsification device was fabricated and used to produce GelMA droplets, following our previous publication [1], which was adopted from an established protocol [2]. Solutions were loaded into 5 mL syringes (BD) and injected into the microfluidic device using two syringe pumps (PHD 2000, Harvard Apparatus) while the flow rates set at 20 and 40 μL $min^{-1}$ for the dispersed and continuous phases, respectively. The microfluidic setup was kept at 37-40°C using a space heater. The droplet suspension in oil and surfactant was then collected into a snap cap 1.7-mL microcentrifuge tube (Globe Scientific) and maintained at 4°C overnight, protected from light, to yield physically crosslinked GelMA microgels.

To form individually photochemically crosslinked microgels, the microgel suspension was centrifuged at 100×*g* for 15 s. Then 100 μL of the suspension was transferred into a snap cap 2-mL microcentrifuge tube (Globe Scientific) using a positive displacement pipette (Microman E M100E, Gilson). The microgels were individually photocrosslinked via exposing to visible light (OmniCure LX500 with 395 nm UV LED head, Excelitas Technologies Corp.) with an intensity = 2 W $cm^{-2}$ for 1 min, vortexed at 3000 rpm using a digital vortex mixer, and exposed to light for an additional 1 min, to form intra-microgel covalent bonds. To remove the oil and surfactant, 100 μL of 20% v/v 1H,1H-perfluoro-1-octanol (PFO, MilliporeSigma) solution in Novec™ 7500 Engineered Fluid was added to the photocrosslinked microgel suspension. The mixture was vortexed at 3000 rpm for 20 s using a digital vortex mixer, followed by centrifugation at 4000×*g* for 15 s. After discarding the oil and surfactant, 200 μL of DPBS was added to the suspension. The mixture was vortexed for 15 s at 3000 rpm, followed by centrifugation at 4000×*g* for 15 s. After removing the remaining oil and surfactant, the process was repeated once until the residual oil and surfactant were completely removed.

To assess microgel stability, the microgel suspension in DPBS was centrifuged at 4000×*g* for 15 s. Then, the DPBS was removed, and 10 μL of packed microgels was pipetted into a Petri dish, containing 5 mL of preheated DPBS, which was placed in an STXG stage-top incubator (Tokai Hit) at 37°C. Images were then acquired from the samples ($n$ = 3) at 5 min intervals for 1 h to monitor microgel stability. Microgel diameter was measured using Fiji ImageJ software (version 1.54f)[3]. The images were converted to 8-bit type, then thresholded, followed by analysis using “Analyze Particle” built-in tool in the software.

**The fabrication of bulk hydrogel scaffolds and rheological characterization**. To measure the mechanical properties of bulk hydrogels, cylindrical GelMA scaffolds were first fabricated and subsequently characterized using oscillatory rheology. To fabricate bulk hydrogel scaffolds, LAP was dissolved in room temperature DPBS , to yield a 0.1% w/v LAP solution. Lyophilized GelMA was then dissolved in the LAP solution to prepare 4 or 15% w/v solutions. The GelMA solutions were then pipetted into laser-cut, disk-shaped molds (diameter = 8 mm and height = 3 mm), made from acrylic sheet (Astra Products), and sandwiched between a microscope slide (bottom) and a cover slip (top). The GelMA-loaded molds were transferred to a custom-built humidity chamber. The chamber was made from a Petri dish (VWR) containing Kimwipes soaked in ultra-pure water. Samples were then maintained at 4 °C overnight, protected from light, to enable physical gel formation. The molded gel was then exposed to visible light (OmniCure LX500 with 405 nm ultraviolet (UV) light-emitting diode (LED) head, Excelitas Technologies) with an intensity = 2 W $cm^{-2}$ for 2 min to photocrosslink the molded GelMA.

The dynamic moduli of bulk hydrogel scaffolds, as estimates for the local rheological properties of microgels, were assessed using oscillatory rheology tests via an AR-G2 rheometer (TA Instruments). To this end, cylindrical scaffolds (diameter = 8 mm and height = 3 mm) were incubated overnight in DPBS at room temperature. Scaffold were then loaded onto the rheometer and sandwiched between parallel plates, while the temperature maintained at 25 °C during the tests. A preliminary strain amplitude sweep test was performed (constant frequency = 1 rad $s^{-1}$) to identify the linear viscoelastic region (LVR). Next, an oscillatory frequency sweep test was conducted within the range of 0.1 to 100 rad $s^{-1}$ (oscillatory strain = 0.1%) ($n$ = 5).

**Measurement of hydraulic conductivity (Lp) in individually perfused rat mesenteric microvessels and VE cadherin microvessel staining.** A single mesenteric microvessel was cannulated with a micropipette and perfused with albumin-Ringer solution (control) containing red blood cells (~1% vol/vol) as markers under a known hydrostatic pressure ranging from 40 to 60 $cmH_2O$. For each measurement, the perfused vessel was occluded briefly downstream with a glass rod for 5~7 s. The initial water flux/unit area of microvessel wall was calculated from the velocity of the marker cell after vessel occlusion, the vessel radius, and the distance between the marker cell and the occlusion site. Lp was calculated as the slope of the relationship between the initial water flow/unit area of vessel wall and the pressure difference across the vessel wall. In each experiment, the baseline Lp and the Lp after application of testing solutions was calculated.

Staining of the perfused vessels were carried out by fixation with 2% paraformaldehyde and permeabilization with .1% Triton X-100 before exposure to VE-Cadherin (Santa Cruz, rabbit) antibody. The tissue was then incubated with Alexa-488 conjugated secondary antibody (Invitrogen, anti-rabbit) at room temperature for 3 h and images were obtained using a Leica SP8 confocal microscope for both images and measurement of fluorescent intensity of vessels per unit area.

**Supplementary Methods References**


1. Ataie, Z., et al., *Gelatin Methacryloyl Granular Hydrogel Scaffolds: High-throughput Microgel Fabrication, Lyophilization, Chemical Assembly, and 3D Bioprinting.* J Vis Exp, 2022(190).
2. de Rutte, J.M., J. Koh, and D. Di Carlo, *Scalable High-Throughput Production of Modular Microgels for In Situ Assembly of Microporous Tissue Scaffolds.* Advanced Functional Materials, 2019. **29**(25): p. 1900071.
3. Schindelin, J., et al., *Fiji: an open-source platform for biological-image analysis.* Nat Methods, 2012. **9**(7): p. 676-82.
